# A PROXIMITY LIGATION SCREEN IDENTIFIES SNAT2 AS A NOVEL TARGET OF THE MARCH1 E3 UBIQUITIN LIGASE

**DOI:** 10.1101/2024.09.16.613264

**Authors:** Renaud Balthazard, William Mitchell, Maxime Raymond, Arnau Ballestero Vidal, Dominic G. Roy, Libia Cecilia Palma Zambrano, Mohamed Abdelwafi Moulefera, Denis Faubert, Sarah Pasquin, Jean-François Gauchat, Jacques Thibodeau

## Abstract

E3 ubiquitin ligases are part of various families of proteins and include hundreds of members, which play key roles in all aspects of cell biology. They generally regulate the half-life of other proteins but can also modulate their cellular localization and functions. The MARCH family of ubiquitin ligases is composed of 11 members and two closely related proteins, MARCH1 and MARCH8, share similar targets, while being active in different cell types. Although they appear to target principally immune cell components, such as MHC class II molecules and the co-stimulatory molecule CD86, the repertory of their targets remains to be fully documented. Here, to further define the MARCH1’s interactome, we adapted a proximity-dependent biotin identification (BioID)-based screening approach in live HEK293 cells. We transfected a fusion protein consisting of mouse MARCH1 linked to YFP at its N-terminus and to the biotin ligase of *Aquifex aeolicus* at its C-terminus. Upon transient overexpression of this construct in the presence of exogenous biotin, we could recover biotinylated proteins that are presumably found within 10nm of MARCH1. To help in the identification of *bona fide* down-regulated specific targets, we compared MARCH1’s interactome with the one obtained using a ubiquitination-deficient MARCH1 mutant (MARCH1W104A). CD98 and CD71, two previously described targets of MARCH1, were identified in this screen. Of 16 other biotinylated proteins identified by semi-quantitative mass spectrometry, 10 were tested directly by flow cytometry to monitor their expression in the presence or absence of transfected MARCH1. The protein levels of five of these endogenous targets, CD29, CD112, NKCC1, CD147 and SNAT2, confirmed their negative regulation by MARCH1 in this system. SNAT2 was particularly sensitive to the presence of MARCH1 and was found to be ubiquitinated on Western blots following immunoprecipitation. Thus, BioID2 is an effective mean of characterizing the interactome of MARCH1 and the identification of SNAT2 suggests a role of this ubiquitin ligase in cellular metabolism.

## INTRODUCTION

Ubiquitination is an important post-translational modification of proteins in eukaryotic cells. It is implicated in a plethora of biological processes, such as protein homeostasis and intracellular transport (Jiang and Chen, 2012; Rodgers et al., 2014; Samji et al., 2014). Ubiquitination is a sequential process involving 3 major steps: first, the E1 protein must activate the ubiquitin, which is then transferred to an E2. These are limited in numbers, and they collaborate with E3 ligases for the conjugation of the ubiquitin moiety to the target. The specificity of the process is inherent to the E3, which recognizes the substrate and catalyzes the covalent transfer of ubiquitin principally to specific lysine residues of the targets (Hershko and Ciechanover, 1998; Hershko et al., 1983). Depending on a number of factors, this first ubiquitin can itself be modified and assembled into long, often branched ubiquitin chains (Pickart and Fushman, 2004).

Membrane associated RING-CH (MARCH) proteins constitute a family of mammalian E3 ubiquitin ligases composed of 11 members (MARCH1-11) (Reviewed in(Bauer et al., 2017). Within the MARCH proteins, phylogenetic analysis identified four groups, which share structural homology and overlapping substrates (Bartee et al., 2004). One of these groups includes MARCH1 and 8, which are multipass proteins with two transmembrane (TM) helical domains also involved in target recognition and dimer formation(Bourgeois-Daigneault and Thibodeau, 2012; Lehner et al., 2005). The TM1 and TM2 are connected via a short luminal loop (LL) of approximately 20 amino acids. Their catalytic RING-CH domain is cytosolic and located at a distance of about 20 residues from a first transmembrane region (TM1) in the N-terminal part of the protein (Reviewed in(Bauer et al., 2017; Deshaies and Joazeiro, 2009). It has been shown that a point mutation at tryptophan 104 in the RING domain of MARCH1 obstructs the interaction with E2 enzymes and drastically decreases the ubiquitin ligase activity (Hewitt et al., 2002; Jabbour et al., 2009; Joazeiro et al., 1999).

MARCH1 is best known for its role in antigen presenting cells (APCs) (Matsuki et al., 2007; Mittal et al., 2015; Ohmura-Hoshino et al., 2006; Walseng et al., 2010). Its expression appears to peak in immature dendritic DCs and resting follicular (FO) B cells (De Gassart et al., 2008; Galbas et al., 2012; Matsuki et al., 2007; Walseng et al., 2010). Still, the half-life of the MARCH1 protein is very short and was estimated to about 30 minutes in primary murine DCs (Jabbour et al., 2009). In resting APCs, MARCH1 ubiquitinates many membrane proteins, thereby preventing their recycling and causing their lysosomal degradation (De Gassart et al., 2008; Thibodeau et al., 2008; Walseng et al., 2010). The best characterized targets of human and mouse MARCH1 are major histocompatibility complex class II molecules (MHCIIs) and CD86. When APCs get activated, MARCH1 is down-regulated to increase the display of peptide-MHCII complexes and co-stimulatory molecules (Cho et al., 2015; De Gassart et al., 2008). Accordingly, DCs and B cells of MARCH1-deficient mice show dramatic expression of MHCIIs and CD86 at their surface (Baravalle et al., 2011; Matsuki et al., 2007), up to a point where the integrity of lipid rafts and tetraspanin domains is compromised (Oh et al., 2018). While the up-regulation of MARCH1 by IL-10 and its homology with subversive viral proteins are in line with immunosuppressive functions (Bartee et al., 2004; Thibodeau et al., 2008), its tight regulation in light and dark zones germinal center (GC) B cells highlights the possible role of this E3 ubiquitin ligase in multiple important physiological processes, such as antibody maturation (Bannard et al., 2016).

Since the discovery of the MARCH proteins, several laboratories have succeeded in identifying specific targets of each family member. For example, techniques like stable isotope labeling by amino acid in cell culture (SILAC), or cell surface proteomic analysis have been adapted to define substrates of many members of the MARCH family, including MARCH1 and MARCH8 (Bartee et al., 2010; Hoer et al., 2007; Sandow et al., 2021; Schriek et al., 2021). Other functional screens have been performed, which revealed the role of MARCH proteins in regulating various receptor-activated pathways. For example, a large screen of about 10000 cDNA clones identified human MARCH8 as a key negative regulator of IL-1β signaling (Chen et al., 2012). Also, using HeLa cells grown in serum-free conditions, a shRNA screen of more than 600 E3 ubiquitin ligases revealed that human MARCH1 controls cell surface insulin receptor levels (Nagarajan et al., 2016). Altogether, these different approaches allowed the identification of novel, potentially biologically relevant MARCH targets. These findings suggest that the effects of MARCH1 may be more diverse than initially anticipated.

To better appreciate the full spectrum of MARCH proteins actions, it will be essential to identify all interacting partners in different cell populations and physiological conditions. Recently, Roux and colleagues developed a method to pin down relevant protein interactions that occur in living cells (Roux et al., 2018; Roux et al., 2012). This technique, called BioID, relies on proximity-dependent biotin labeling and requires the protein of interest to be fused to a constitutively active biotin-conjugating enzyme of bacterial origin. The biotinoyl-AMPs that it releases can thus simply diffuse and react with the surrounding amine groups, including those present on the lysine residues of neighboring polypeptides (Roux et al., 2018; Roux et al., 2012). Because biotinylation is a rare protein modification in nature and isn’t normally found in eukaryotes, the modified proteins can first be easily and specifically captured using avidin beads, then identified by mass spectrometry (Coyaud et al., 2015; Roux et al., 2018). Therefore, BioID appears to be a promising approach to catch the breadth of the different MARCH protein networks.

Here, we aimed to further characterize the MARCH1’s interactome by using BioID in HEK293 cells. While we found some previously described MARCH1 targets, such as transferrin receptor (CD71), we also uncovered some new ones, including the amino acid transporter SNAT2 (SLC38A2). Our results point to new potential molecular handles through which MARCH1 can control immune responses and highlight the usefulness of BioID for the study of MARCH proteins.

## MATERIAL and METHODS

### Plasmids

The small and efficient BioID2 biotin ligase from the gram-negative bacterium *Aquifex aeolicus* (Kim et al., 2016) was fused to the C-terminal end of mouse MARCH1 (mMARCH1). The enhanced yellow fluorescent protein (hereafter YFP) was attached at the N-terminus of mMARCH1. The DNA coding for this fusion protein (YFP-MARCH1-BioID2) was codon-optimized, synthetized and cloned in the pcDNA3.4 expression vector (GeneArt, ThermoFisher scientific, Whitby, Ontario). The complete nucleotide sequence is shown in Figure S1A. To replace MARCH1 with other proteins of interest, restriction sites were introduced between YFP and mMARCH1, and between mMARCH1 and BioID2 coding sequences. The labeling radius of BioID2 is estimated to be about 10 to 15 nm (Kim et al., 2016) but this range can be increased by the inclusion of a flexible linker. To do so, we introduced a *BstX*I restriction site between the MARCH1 and BioID2 coding sequences. Also, this restriction site was designed to be compatible with the 3’ overhang sequence of KpnI-cut DNA, thereby allowing the removal of MARCH1 and, upon religation, the making of a control fusion protein devoid of any ubiquitin ligase (YFP-BioID2). The YFP-MARCH1W104A-BioID2 plasmid, encoding the inactive E3 ubiquitin ligase, was generated from YFP-MARCH1-BioID2 by site-directed mutagenesis using the PCR overlap extension method (Ho et al., 1989). Plasmids pcDNA3.1 encoding the human MARCH constructs YFP-MARCH1, YFP-MARCH1W104I, YFP-MARCH8 or GFP-MARCH9 have been described previously(Bourgeois-Daigneault and Thibodeau, 2012). The codon-optimized DNAs coding for the SNAT2-FLAG and SNAT2 7KR-FLAG fusion proteins, cloned in pcDNA3.1, were purchased from GenScript (Piscataway, NJ). The nucleotide sequences are shown in Figure S1B.

#### Cell culture and transfections

HEK293 cells were cultured in DMEM supplemented with 5% FBS (Wisent, St-Bruno, Quebec) and Glutamax (Gibco, ThermoFisher scientific, Whitby, Ontario). Cells were transiently transfected with polyethyleneimine, as previously described (Cloutier et al., 2014).

#### siRNA

siRNA SLC38A2 (s634) Silencer Select and Negative Control No. 2 (Invitrogen, ThermoFisher scientific) were transfected in HEK293 cells using Lipofectamine RNAiMax (Invitrogen), following the manufacturer’s protocol.

#### Flow Cytometry

Cells were fixed in 4% paraformaldehyde and permeabilized in PBS containing 0.1% saponin and 1% BSA on ice. Cells were stained in permeabilization buffer with the indicated antibodies and analyzed on a FACSCanto II or FACSymphony A1 (BD, Mississauga, Ontario). Graphs were prepared using FlowJo.

#### Western blots

Cells were lysed for 30 min on ice in 1% Triton X-100 buffer containing Mg132 (10µM), N-Ethylmaleimide (0.2mM) and protease inhibitors (Sigma, Oakville, Ontario). Post nuclear supernatants were prepared and proteins were denatured in Laemmli buffer. After boiling, proteins were resolved by SDS-PAGE and transferred to a nitrocellulose membrane. After blocking in reconstituted powdered milk, primary antibodies were incubated overnight at 4°C. HRP-coupled anti-ubiquitin P4D1 antibody (Biolegend, San-Diego, CA) was incubated at room temperature for 2 hours. Nitrocellulose membranes were revealed with ECL solution (Amersham, Little Chalfont, UK) on an Amersham Imager 600.

#### Antibodies and Reagents

Antibodies against CD71 (Clone CY1G4), FLAG (L5), CD29 (F2/16), CD112 (TX31), as well as Alexa fluor 647-coupled Goat anti-Mouse (AF647; Poly4053) and APC-coupled Streptavidin were purchased from Biolegend (San-Diego, CA). Antibodies specific for SNAT2 (C-6), CD147 (8D6), NKCC1 (F-4), AE2 (D-3), MARK3 (F-6), Occludin (E-5) and SNAT1 (H-9) were purchased from Santa Cruz Biotechnology (Dallas, TX).

#### BioID2

HEK293 cells were transfected with the indicated plasmids. After 24 hours, cells were activated with Sendai virus (Cantell strain; a gift from Dr. Daniel Lamarre, IRIC, Montreal) and biotin was added at a final concentration of 50µM for an additional 16 hours (Liu et al., 2020). Cells were frozen at -80°C and lysed in cold RIPA buffer supplemented with proteases inhibitors. In each sample, 250 units of benzonases (Sigma, Oakville, Ontario) was added. Samples were sonicated (3x 10s at 30% amplitude) on ice. Supernatants were cleared by centrifugation at 13 000 rpm for 30 minutes at 4°C. Then, 70ul of streptavidin-sepharose slurry (GE Healthcare, Bensalem, PA) was added in each sample. After 4 hours with rotation, the beads were washed in RIPA buffer and then in ammonium bicarbonate 50mM at 4°C. Beads were resuspended in ammonium bicarbonate and proteins digested on the same day using 0.5 ug Sequencing Grade Modified Trypsin (Promega, Madison, WI) overnight at 37°C with agitation. The supernatants were collected, and the beads were washed twice with 100 uL water. The supernatants of each wash were pooled, reduced and alkylated. The reduction step is done with 9 mM dithiothreitol at 37°C for 30 minutes and, after cooling for 10 minutes, the alkylation step is done with 17 mM iodoacetamide at room temperature for 20 minutes in the dark. The supernatants were acidified with trifluoroacetic acid for desalting and removal of residual detergents by MCX (Waters Oasis MCX 96-well Elution Plate) following the manufacturer’s instructions. After elution in 10% ammonium hydroxide /90% methanol (v/v), samples were dried with a Speed-vac, reconstituted under agitation for 15 min in 15 µL of 5% FA and loaded into a 75 μm i.d. × 150 mm Self-Pack C18 column installed in the Easy-nLC II system (Proxeon Biosystems, Thermo Fisher Scientific, Whitby, Ontario). The buffers used for chromatography were 0.2% formic acid (buffer A) and 90% acetonitrile/0.2% formic acid (buffer B). Peptides were eluted with a two-slope gradient at a flowrate of 250 nL/min. Solvent B first increased from 1 to 35% in 105 min and then from 35 to 84% B in 25 min. The HPLC system was coupled to a Q Exactive mass spectrometer (Thermo Fisher Scientific, Whitby, Ontario) through a Nanospray Flex Ion Source. Nanospray and S-lens voltages were set to 1.3-1.8 kV and 50 V, respectively. Capillary temperature was set to 225 °C. Full scan mass spectrometry (MS) survey spectra (m/z 360-2000) in profile mode were acquired in the Orbitrap with a resolution of 70 000. The 16 most intense peptide ions were fragmented in the collision cell and MS/MS spectra were analyzed in the Orbitrap.

#### Protein identification and data filtration

The peak list files were generated with Proteome Discoverer (version 2.1) using the following parameters: minimum mass set to 500 Da, maximum mass set to 6000 Da, no grouping of MS/MS spectra, precursor charge set to auto, and minimum number of fragment ions set to 5. Protein database searching was performed with Mascot 2.6 (Matrix Science, Boston, MA) against the Refseq human protein database (May 16^th^, 2018). The mass tolerances for precursor and fragment ions were set to 10 ppm and 0.6 Da, respectively. Trypsin was used as the enzyme allowing for up to 1 missed cleavage. Cysteine carbamidomethylation was specified as a fixed modification, and methionine oxidation as variable modifications. Data interpretation was performed using Scaffold (version 4.8). After data filtration, a node graph of the remaining proteins was created using Cytoscape v3.10 and stringApp2.0 for integration to the STRING database of protein interactions. A STRING score of 0.3 was used to define meaningful partnership (Doncheva et al., 2023; Szklarczyk et al., 2023)

## RESULTS

The biotin-conjugating enzyme originally used in standard BioID protocols was an R118G mutant of *Escherichia coli*’s BirA. It allowed the characterization of the interactome of many proteins, including different types of E3 ubiquitin ligases (Coyaud et al., 2015; Dho et al., 2019). However, an important challenge for any method that relies on the expression of fusion constructs is to ensure proper folding, biological activity and subcellular targeting of the chimeric protein. Although BirA (321 amino acids) is only slightly larger than the green fluorescent protein (GFP) (239 amino acids) successfully linked previously to MARCH proteins, its presence may affect the properties of certain constructs. Therefore, to minimize the impact of the biotin ligase moiety, we used a smaller enzyme isolated from the bacterium *Aquifex aeolicus* and recently introduced by the group of Roux. This method, called BioID2, allows more selective targeting of fusion proteins and exhibits improved labeling of neighboring interactors(Kim et al., 2016).

### YFP-MARCH-BioID2 is capable of ubiquitination and biotinylation

To further characterize the interactome of MARCH1, we used the BioID2 proximity ligation approach. Including YFP in the construct facilitates the tracking and quantification of MARCH1 and BioID2 in transfected cells. Previous studies in our laboratory showed that a fusion protein made of MARCH1 with an YFP moiety at its N-terminus maintained its ability to ubiquitinate and regulate the surface expression of MHCII molecules (Bourgeois-Daigneault and Thibodeau, 2012). Thus, we opted for a construct in which MARCH1 is sandwiched between YFP at its N-terminus and the biotin ligase (hereafter called BioID2) at its C-terminus (YFP-MARCH1-BioID2; Figure 1A). To control for the non-specific MARCH1-independent biotinylation of intracellular proteins, we generated another fusion protein consisting only of YFP and the biotin ligase (YFP-BioID2). The cell surface down-regulation of CD71, a known target of MARCH1(Bartee et al., 2006), confirmed the ubiquitin ligase activity of the HEK293-transfected YFP-MARCH1-BioID2 fusion protein (Figure 1B).

**Figure 1.**
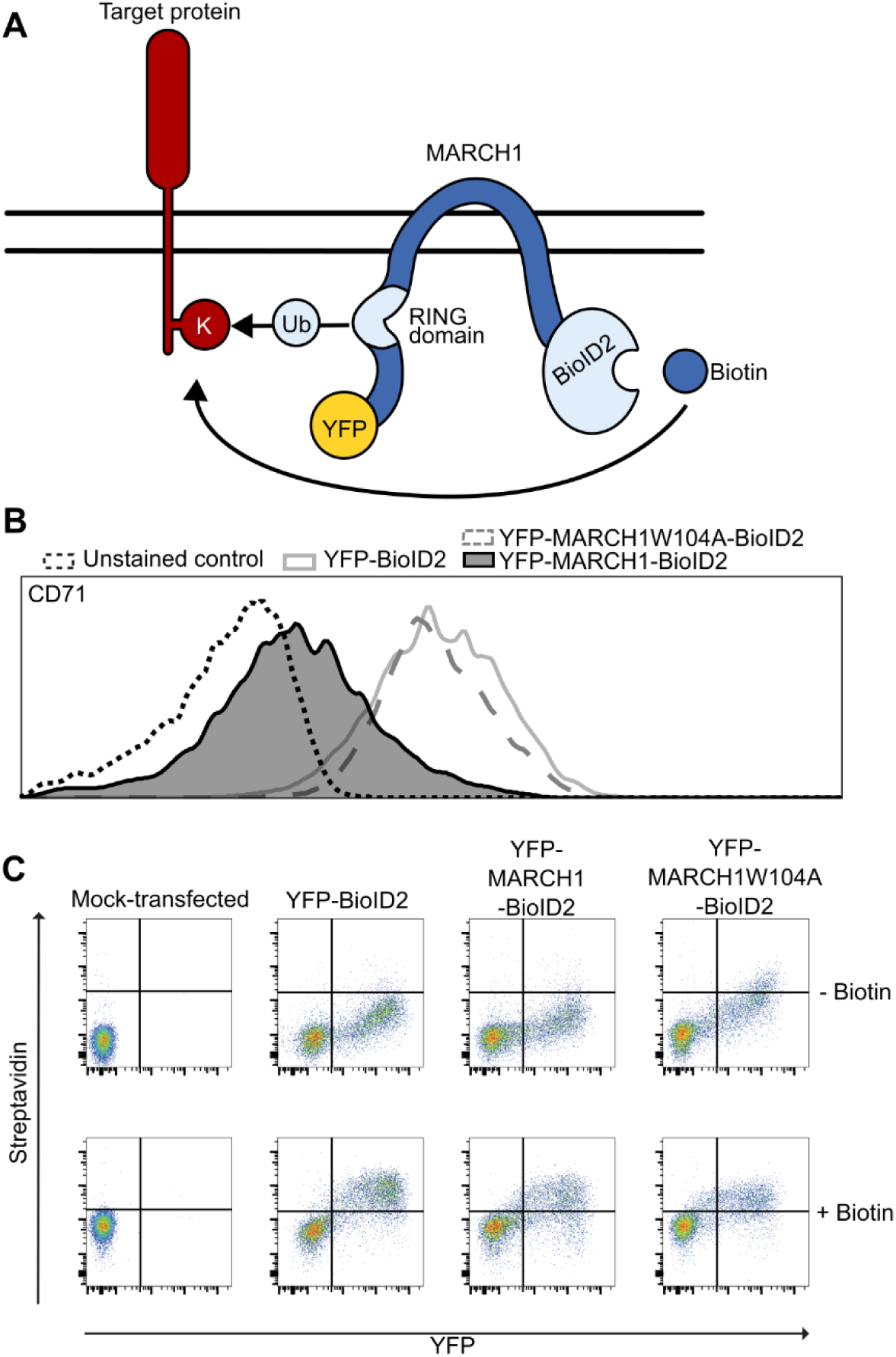
The YFP-MARCH1-BioID2 fusion protein conserves both ubiquitin ligase and biotin ligase activities. **A.** Schematic representation of the recombinant enzyme. YFP was fused in frame to the N-terminus of mouse MARCH1. The biotin ligase of *Aquifex aeolicus* (BioID2) was fused to the C-terminus of YFP-MARCH1. **B.** Fusion proteins (YFP-BioID2, YFP-MARCH1-BioID2, and YFP-MARCH1W104A-BioID2) were transiently expressed separately in HEK293 cells for 48h and the surface expression of CD71 was monitored by flow cytometry. **C.** HEK293 cells were transfected with the plasmids coding for YFP-BioID2, YFP-MARCH1-BioID2, or YFP-MARCH1W104A-BioID2. After 24 hours, biotin was added to the culture medium at a final concentration of 50 µM for 16 hours. Control cells (top row) did not receive biotin. Cells were then permeabilized, washed, stained with fluorescent streptavidin, and analyzed by flow cytometry.

We then verified that BioID2 was functional when fused at its N-terminus to YFP-MARCH1. Biotinylation was assessed by intracellular staining of transiently transfected HEK293 cells using fluorescent avidin. Figure 1C shows that upon addition of exogenous biotin to live cells, a strong positive correlation was observed between the avidin signal and the presence of BioID2, which is monitored by the YFP signal.

### BioID2 identifies new MARCH1 targets

The YFP-BioID2 and YFP-MARCH1-BioID2 fusion proteins were transiently overexpressed, in triplicates, in HEK293 cells. Biotin was added and cells were kept in culture for another 16h before lysis. To increase the yield of biotinylated proteins, we also used a mutated MARCH1 (YFP-MARCH1W104A-BioID2), which is greatly impaired in its ability to interact with E2s and to ubiquitinate its targets (Jabbour et al., 2009). Such a strategy, based on the comparison between the active and inactive forms of an E3 ubiquitin ligase, has been used by others in the past, for example as part of label-free shotgun proteomics (Burande et al., 2009). Proteins that are underrepresented in the MARCH1 samples as compared to MARCH1W104A are most likely direct targets of this ubiquitin ligase. The proteins equally represented in both samples are believed to come in the vicinity of MARCH1 without being ubiquitinated nor degraded. In our system, YFP-MARCH1W104A-BioID2 was indeed practically inactive and unable to down-regulate CD71 (Figure 1B).

Biotinylated material was extracted from the cell lysates using avidin beads. Proteins were digested and analyzed by MS. All conditions combined, a total of 3187 proteins were identified (Fig. 2A and S2). Then, proteins were disregarded if not represented in all replicates of both YFP-MARCH1-BioID2 and MARCH1W104A-BioID2 samples. Of the remaining 1819 putative targets of MARCH1, only 84 proteins showed a statistically significant (p<0.05 with one-tailed Welch corrected T-test) reduction (> 2-fold) in spectral counts for YFP-MARCH1-BioID2, as compared to MARCH1W104A-BioID2. This cutoff, as mentioned above, skews the results toward proteins that are most likely ubiquitinated by MARCH1, and possibly degraded. Of the 84 likely interactors, 55 had an average hit count below 10 for the control YFP-BioID2. We further shortened the list of potential MARCH1 targets 1) by removing any protein that was identified in all 3 replicates of the negative control YFP-BioID2, and 2) by excluding less abundant proteins showing an average YFP-MARCH1W104A-BioID2 spectral count below 9. Table I lists the remaining 18 proteins, which include CD71 and CD98, two well-described targets of MARCH1(Ablack et al., 2015; Fujita et al., 2013) Figure 2B shows a connectivity map based on known protein-protein interactions from the STRING database. (Szklarczyk et al., 2023). Thirteen of these targets have been shown, as expected, to be transmembrane proteins that can be detected at the plasma membrane, and which cytoplasmic domains contain at least one lysine residue (Table SI). MARK3, MARCKS and SNAP23 do not appear to contain transmembrane domains but have been found in association with the inner leaflet of the plasma membrane via binding to phospholipids, protein kinase C and other SNARE proteins, respectively (Hatsuzawa and Sakurai, 2020; Moravcevic et al., 2010; Thelen et al., 1991). In the case of AHCYL and SPTBN2, there is compelling evidence that the proteins can be found associated with the plasma membrane, but the mechanisms remain poorly characterized (Ando et al., 2009; Deng et al., 2024). Nevertheless, all 18 proteins on the short list contain one or more lysine residues, making them valid targets for biotinylation by MARCH1-BioID2.

**Figure 2.**
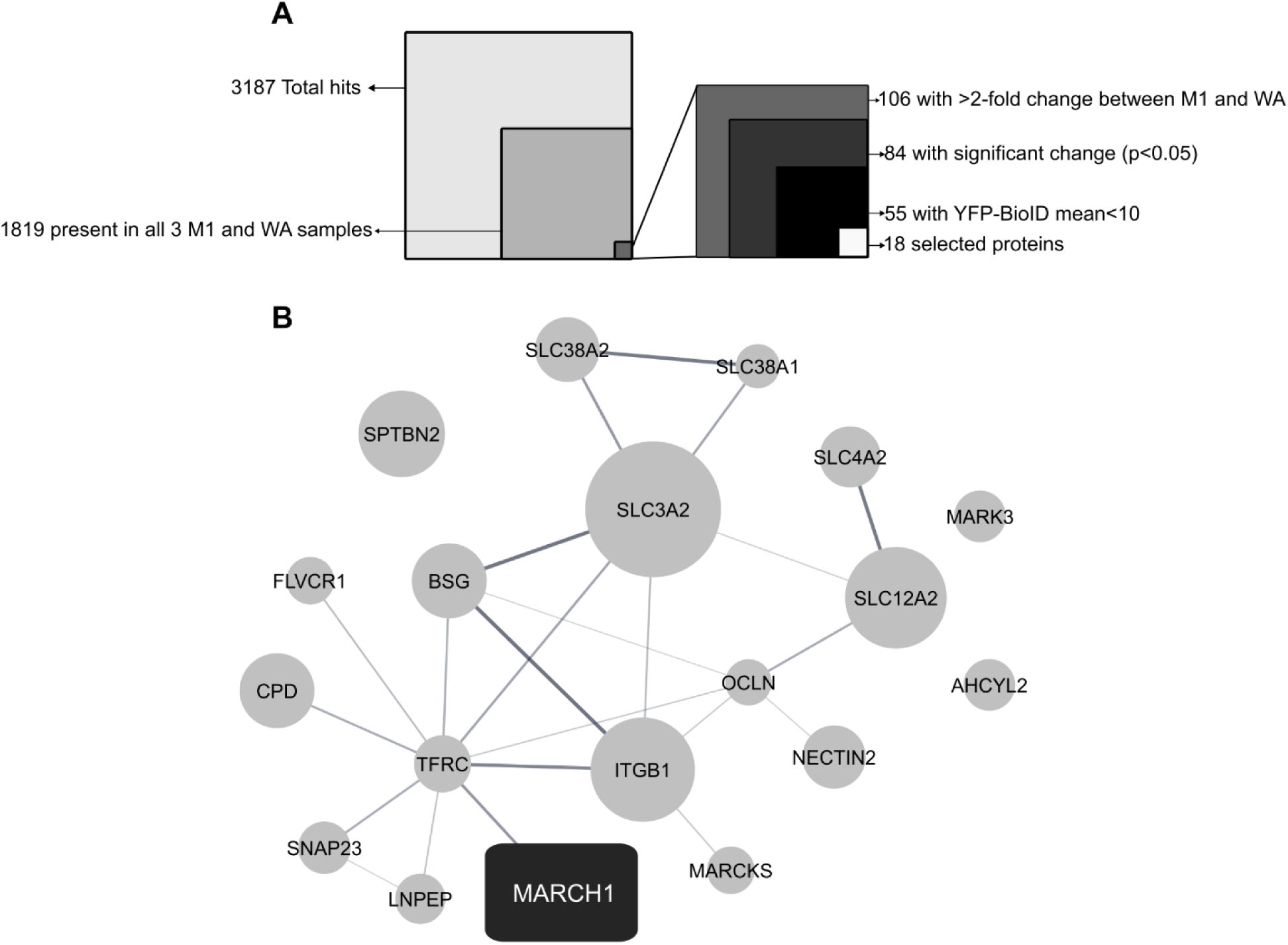
Identification of proteins interacting with MARCH1. YFP-MARCH1-BioID2, YFP-MARCH1W104A-BioID2 and YFP-BioID2 were transiently expressed in HEK293 cells in triplicates. Biotinylated proteins were recovered with avidin-couple beads and analyzed by mass spectrometry. **A.** Breakdown of the hits leading to the final list of interacting proteins (see text for details). **B.** Connectivity map of interactions between the 18 proteins identified (see Table I). STRING database interaction score is represented by line darkness. Node size indicates average protein expression in YFP-MARCH1W104A-BioID2 samples.

Out of the 16 new putative MARCH1 targets (Table I), commercial antibodies validated for flow cytometry were readily available for 10 of them. To directly assess if MARCH1 affects their level of expression, we transiently expressed YFP-MARCH1 into HEK293 and monitored protein levels by flow cytometry in permeabilized cells. By comparing the signal between YFP-negative and YFP-positive cells for each antibody, we concluded that the expression of SNAT2, NKCC1 and CD147 was strongly negatively regulated in MARCH1-expressing cells, while SNAT1, CD29 and CD112 were slightly down-regulated. (Fig. 3). The stronger expression of some of these targets in cells expressing the YFP-MARCH1W104I inactive mutant confirmed that the ubiquitin ligase activity of MARCH1 is responsible for the observed down-regulation (Fig. 4).

**Figure 3.**
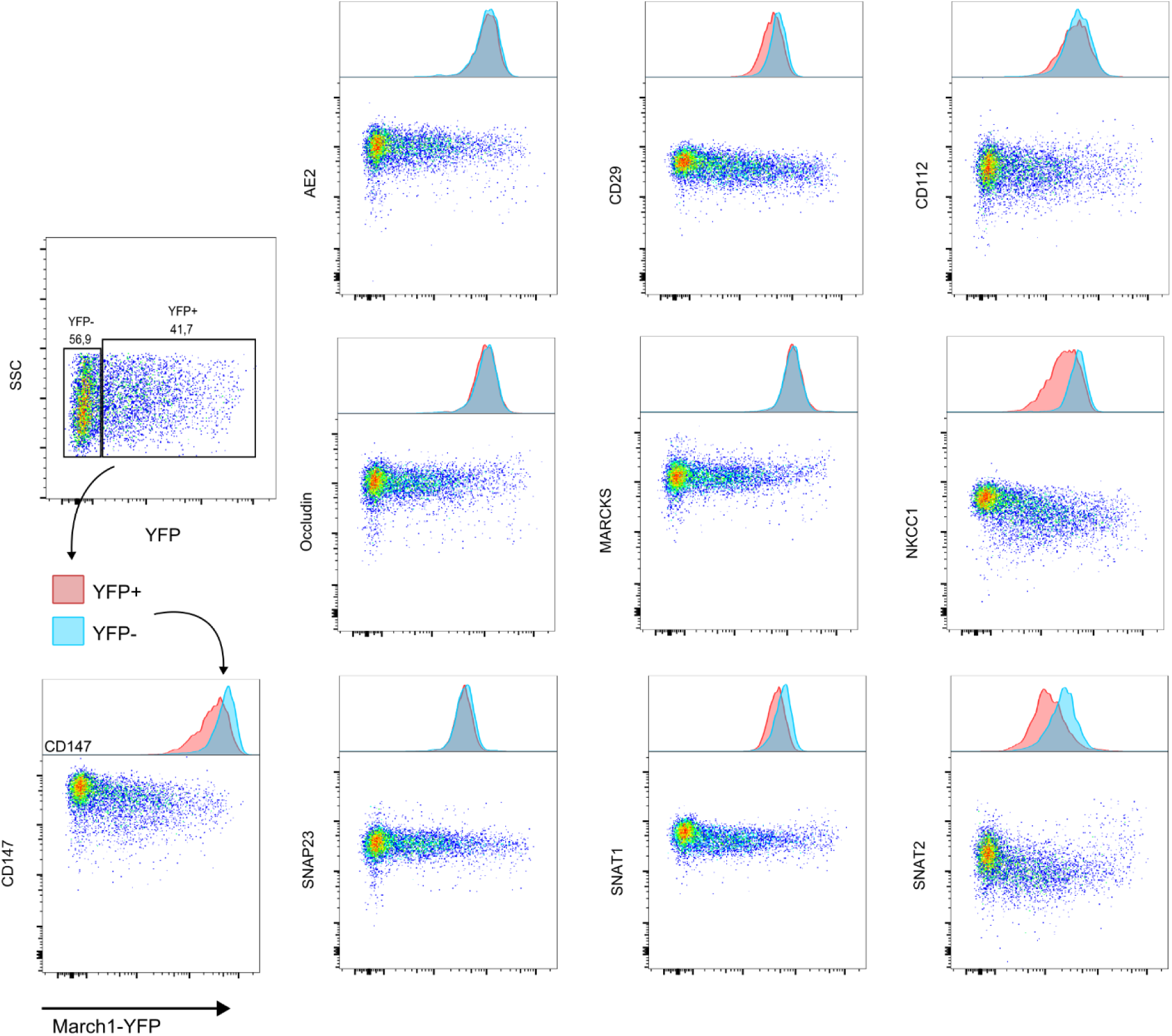
YFP-MARCH1 downregulates proteins levels of several putative endogenous interacting partners. HEK293 cells were transfected with human YFP-MARCH1. After 48h, the cells were permeabilized and analyzed by flow cytometry for the intracellular expression of AE2 (SLC4A2), ITGB1 (CD29), Nectin-2 (CD112), BASIGIN (CD147), MARCKS, NKCC1 (SCL12A1), Occludin, SNAP23, SNAT1 (SLC38A1) and SNAT2 (SLC38A2), which were all identified as potential MARCH1 targets. Histograms over each dot-plot highlight the differential expression of target proteins along the presence or absence of YFP-MARCH1.

**Figure 4.**
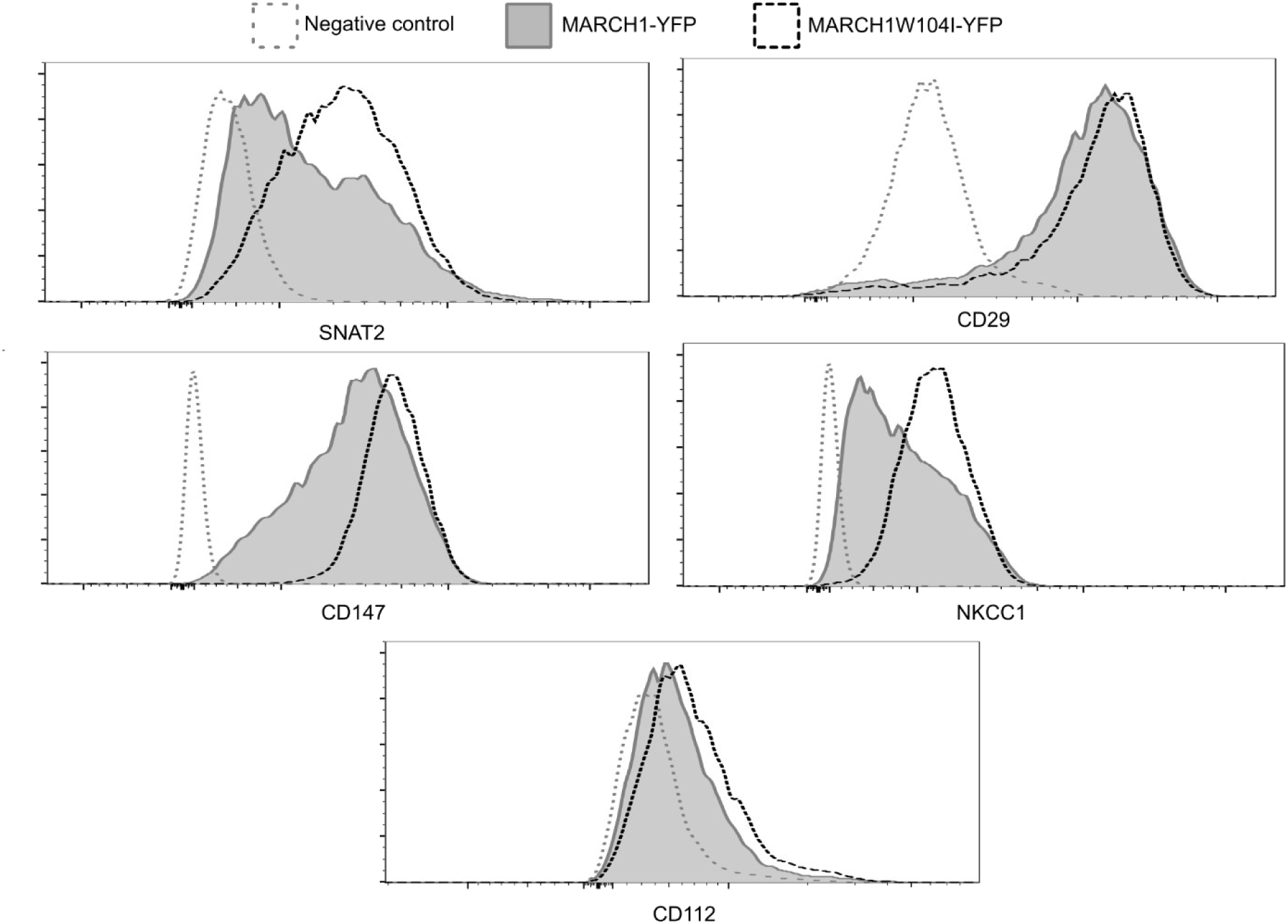
The down-regulation of endogenous targets necessitates the ubiquitination activity of MARCH1. HEK293 cells were transfected with human YFP-MARCH1 or YFP-MARCH1W104I and analyzed 48 hours later for the intracellular expression of SNAT2, CD147, CD29, CD112 and NKCC1 in YFP-expressing cells.

### SNAT2 is ubiquitinated by MARCH1

SNAT2 was the one target showing the most pronounced overall protein down-regulation in the above-described flow cytometry experiments using permeabilized cells. The antibody used in these experiments is highly specific as a siRNA knocking down SNAT2 strongly reduced the signal obtained by flow cytometry (Fig. S3). Since ubiquitination by MARCH1 usually results in diminished cell surface expression of targets, we next determined the impact of this ubiquitin ligase on the cell surface expression of SNAT2. To do so, we transiently transfected YFP-MARCH1 and YFP-MARCH1W104I, along with a tagged SNAT2 in HEK293 cells. The latter has an extracellular C-terminal FLAG epitope (SNAT2-FLAG) that allows for the cell surface staining of the protein (Fig. 5A). Our data show that in the presence of WT MARCH1, SNAT2 expression was markedly reduced at the cell surface (Fig. 5B, left panel). CD29 and CD112 were also found to be down-regulated from the plasma membrane in the presence of MARCH1 (Fig. S4). SNAT2-FLAG was immunoprecipitated with the FLAG-specific antibody and analyzed by Western blotting for the presence of ubiquitin. Figure 5C shows that the ubiquitin-specific P4D1 mAb produced a strong smear of ubiquitinated material when SNAT2-FLAG was co-expressed with MARCH1, but not when expressed alone or together with MARCH1W104I.

**Figure 5.**
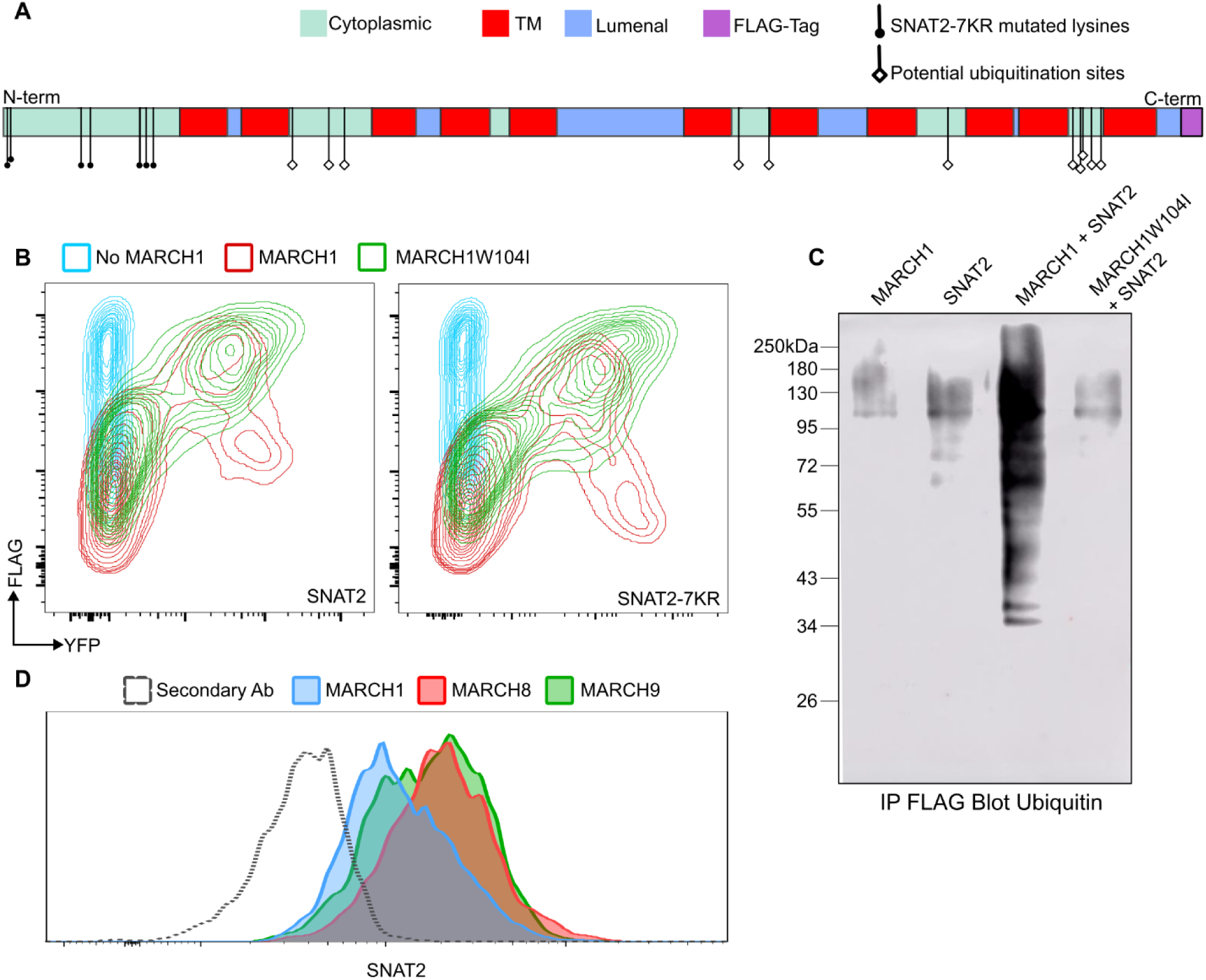
SNAT2 is ubiquitinated and down-regulated from the plasma membrane by MARCH1. **A.** Map of the SNAT2-FLAG protein showing its cytoplasmic regions in green, the transmembrane segments in red and external domains in blue. The FLAG tag is represented in purple. The N-terminal lysines mutated for arginines in SNAT2-7KR-FLAG are shown as black circles while other internal lysines that could potentially be ubiquitinated by MARCH1 are shown as white diamonds. **B.** HEK293 cells were transfected with SNAT2-FLAG or SNAT2-7KR-FLAG alone, or in combination with YFP-MARCH1 and YFP-MARCH1W104I. After 48h, cell surface expression of SNAT2 was measured by flow cytometry using the anti-FLAG antibody. **C.** HEK293 cells were transfected with SNAT2-FLAG alone, or together with YFP-MARCH1 or YFP-MARCH1W104I. YFP-MARCH1 was also transfected alone as an additional control. Cells were lysed and proteins immunoprecipitated using a FLAG-specific antibody coupled to protein-G sepharose beads. Proteins were analyzed by Western blotting using the ubiquitin-specific P4D1 antibody directly coupled to HRP. **D.** HEK293 cells were transfected with human YFP-MARCH1, YFP-MARCH8 or GFP-MARCH9. After 48h, cells were permeabilized and analyzed by flow cytometry for the expression of SNAT2 in YFP/GFP+ gated cells.

A mutated SNAT2-FLAG with its first seven N-terminal lysine residues swapped for arginine (SNAT2-7KR-FLAG; Fig. 5A) was transfected in HEK293 cells along with MARCH1 or MARCH1W104I. After 48 hours, SNAT2 expression was measured by flow cytometry using the FLAG-specific antibody. Despite the removal of the N-terminal cytoplasmic domain lysines, which we hypothesized would encompass the most likely ubiquitinated residue, SNAT2-7KR-FLAG was very efficiently down regulated by MARCH1 (Fig. 5B, right panel). If not on non-lysine amino acids of the N-terminal region, SNAT2 ubiquitination could therefore occur on lysine residues exposed to MARCH1 on the intracytoplasmic loops of the protein, of which we count 11 (Fig. 5A).

In light of the redundancy often observed between MARCH proteins, for example in the recognition of MHC class II molecules by both MARCH1 and MARCH8 (Matsuki et al., 2007), we tested other members of the family for their capacity to downregulate endogenous SNAT2. Figure 5D reveals that while overexpressed MARCH8 had no impact over the expression of SNAT2, MARCH9 did have an effect. These results confirm the specificity of the interaction between SNAT2 and MARCH1, but also suggest that other members of the MARCH family could target this transporter.

## DISCUSSION

The goal of the experimental approach described here was to deliver molecular tools that would allow for the screening of physiologically relevant proximity partners of MARCH1 in cell lines and, ultimately, primary mouse and human cells. Our proof-of-concept experiments in HEK293 cells clearly demonstrate the usefulness of the approach. The fusion of BioID2 to the closely related members of the MARCH family, such as MARCH1 and MARCH8, could yield interesting insights into their specific interacting networks. Indeed, using a biotin-based proximity ligation assay, ARFGAP1 was recently identified as a new target of MARCH5 (Barroso-Gomila et al., 2023).

While the design proposed here for the MARCH1 fusion protein permitted the identification of a few thousands potential interacting partners, it is predicted that fusing BioID2 to the N-terminus of MARCH1 instead of the C-terminus would yield some new interactors. In fact, one can think of many other ways of designing the fusion proteins; for example, the YFP moiety could be omitted, and serine-glycine linkers of various lengths could be tested to allow the biotin ligase to move more freely. Interestingly, variants of the biotin ligase have been introduced over the years. TurboID, which relies on an even smaller and more efficient biotin ligase, requires only 30 min to biotinylate substantial amounts of proteins (Branon et al., 2018; Kim et al., 2016). This could be particularly useful for proteins with short half-life or to identify more transient protein-protein interactions. Other biotin ligases, such as UltraID and AirID, have been developed with the goal of increasing specificity as well as reducing background and toxicity (Kido et al., 2020; Kubitz et al., 2022). In addition to the structure/function considerations inherent to the configuration of the fusion proteins, there is also some latitude in the bioinformatic analysis of the results. The various cut-offs and controls used to determine if a given protein is a genuine interactor, or if it should be considered as background, can vary.

Our approach is semi-quantitative but the comparison between the ubiquitination-proficient YFP-MARCH1-BioID and the ubiquitination-deficient YFP-MARCH1W104A-BioID is predicted to provide some interesting insights into the nature of the proteins that were identified. In the analysis presented here, the filters and cut-offs used aimed at identifying targets that are most likely ubiquitinated, excluding proteins merely located close to MARCH1 or interacting without being ubiquitinated, as would be expected of E2s, for example. Thus, it was somewhat surprising to find that a protein, such as occludin, was less represented in the MS analysis with MARCH1 than with MARCH1W104A but did not appear to be degraded by MARCH1 when permeabilized cells were analyzed by flow cytometry. This might be due to the fact that our indirect staining protocol produces a strong background that masks subtle differences. It is also possible that some ubiquitinated targets are simply not degraded but redistributed in early, less degradative endosomes. In support of this, our lab has previously shown that MHC class II molecules were efficiently internalized from the plasma membrane by MARCH1 in HEK293 and Hela cells but were degraded only in the latter (Gauvreau et al., 2009). This is reminiscent of results published by the group of Mellman showing that the length of ubiquitin chains produced by MARCH1, as well as the fate of the modified targets, differ between B cells and DCs (Ma et al., 2012). Altogether, these results highlight the fact that the lack of an effect of MARCH1 on a given protein in one cell type should not be generalized to cells of all origins. Also, it would be useful to test and develop new proximity ligation protocols designed specifically for the identification of ubiquitin ligases’ substrates. For example, BioE3 makes use of a ubiquitin moiety that comprises a low affinity AviTag, thereby increasing the recovery of biotinylated proteins that are genuine targets of the E3 ligase of interest (Barroso-Gomila et al., 2023).

Recently, Villadangos and collaborators conducted a systematic and careful proteomic analysis of the plasma membrane from WT and MARCH1-deficient primary mouse B cells and DCs (Schriek et al., 2021).They concluded that overall, MARCH1 is primarily functional in hematopoietic cells and specifically targets MHC class II molecules and CD86. While we acknowledge that MARCH1 may not be normally expressed in epithelial cells (such as HEK293), and if so, certainly not at the high levels achieved here following transient transfection, it is nevertheless important to document any occurring interaction as the results, be it in cell lines, may ultimately prove to be physiologically important in certain conditions, cell types or tissues (Bandola-Simon and Roche, 2023). For example, Nagarajan et al. have adopted a large-scale RNA interference approach to down-regulate more than 600 known E3 ubiquitin ligases in HeLa cells, with the goal of identifying those that would favor growth in serum-free medium supplemented with sub-optimal insulin concentrations (Nagarajan et al., 2016). Surprisingly, the downregulation of MARCH1 conferred an insulin-sensitizing effect, allowing the growth of these altered HeLa cells. MARCH1 ubiquitinates the insulin receptor and, interestingly, the expression of this E3 ubiquitin ligase is increased in obese white adipose tissues of mice and humans (Nagarajan et al., 2016). Furthermore, while the MARCH1-Tfr interaction was initially mapped in an overexpressing cell line model(Bartee et al., 2004), it has been shown recently that MARCH1 is induced by CMV and regulates iron levels to increase viral replication (Martin et al., 2022). In HIV-infected cells, MARCH1 acts as a restriction factor and down-regulates surface envelope proteins to limit viral infectivity (Zhang et al., 2019). As research on MARCH1/8 progresses, it will become clear that their impact extends beyond immune cells and the downregulation of MHCII or CD86.

Our experiments have led to the identification of new potential targets of MARCH1. If ubiquitinated, it is reasonable to think that proteins such as AHCYL2, SNAP23, MARK3, MARCKS and SPTBN2 are included into multi-protein complexes. Indeed, they lack a TM domain, which was shown to mediate the interaction with MARCH1 (Bourgeois-Daigneault and Thibodeau, 2012; Jabbour et al., 2009). Still, it appears that these proteins can all associate at some point with the plasma membrane, in accordance with a role of MARCH1 at this subcellular location or in recycling endosomes (Ando et al., 2009; Deng et al., 2024; Moravcevic et al., 2010; Naskar and Puri, 2017; Thelen et al., 1991). Interestingly, AHCYL2 (also known as Long-IRBIT) was recently shown to inhibit the lysosomal degradation of AE2 (SLC4A2), an essential anion-exchange transmembrane protein that was also identified here as a MARCH1 target (Itoh et al., 2021). Furthermore, the identification of SNAP23 could be highly relevant given its role in phagosome biogenesis and maturation, as well as in cross-presentation by dendritic cells (Nair-Gupta et al., 2014; Sakurai et al., 2012). MARK3 can interact with microtubule-associated proteins and binds acidic phospholipids in cellular membranes, increasing the likelihood of encountering MARCH1 (Moravcevic et al., 2010; Yang et al., 2023). MARCKS is particularly interesting in light of its association to the inner leaflet of the plasma membrane and its ability to regulate various processes, such as TLR4 signaling and phagocytosis (Carballo et al., 1999; Issara-Amphorn et al., 2023; Thelen et al., 1991). Finally, SPTBN2 (also known as Spectrin beta III) was recently shown to reduce ferroptosis in cancer cells by fine tuning the subcellular localization of SLC7A11 (Deng et al., 2024). Since SLC7A11 and CD98 interact to control intake of cystine and export of glutamate, it appears as if MARCH1 could modulate the trafficking of the complex by interacting with multiple subunits (Li et al., 2022). Exciting results have recently shown that SLC7A11 negatively impacts the capacity of skin DCs to mediate efferocytosis (Maschalidi et al., 2022), suggesting that MARCH1 could have an important regulatory role in this process. However, we do not know if SPTBN2 is actually ubiquitinated, but it comes close enough to MARCH1-BioID2 to be biotinylated. The same applies to CD147 (Basigin) and CD29 (ITGB1), which also associate with CD98hc (Kim and Cho, 2016), In the case of these two proteins however, they are clearly down-regulated by MARCH1 in our system, suggesting roles in various physiological processes, including cell adhesion and cancer (Nyalali et al., 2023; Winograd-Katz et al., 2014). Future studies should determine if they are ubiquitinated or if they are dragged to degradative compartments because of their association with ubiquitinated CD98hc. Of note, others have reported that the effect of MARCH1 on CD147 was marginal in HeLa cells (Eyster et al., 2011) while CD29 was recently shown to be a target of MARCH2. (Sandow et al., 2021).

Three other putative partners (FLVCR1, LNPEP, CPD) containing transmembrane domains have been identified, but were not further characterized here. Their functions remain under investigation by different groups but based on current knowledge, it is likely that they play a role in immunity and that they could serve as MARCH1 targets. Briefly, FLVCR1 (or SLC49A1) is an essential plasma membrane choline transporter and deletion of its genes affects the integrity of mitochondria (Fiorito and Tolosano, 2024; Kenny et al., 2023). The leucyl-cystinyl aminopeptidase LNPEP (also known as IRAP, VP165, AT4R and other names) is a type II integral membrane protein that labels endosomes implicated in cross-presentation by MHC class I molecules (Keller et al., 1995; Saveanu et al., 2009). Carboxypeptidase D (CPD) is a type I transmembrane protein that recycles between the trans-Golgi and the plasma membrane (Varlamov and Fricker, 1998). Among other things, it was shown to regulate nitric oxide synthesis in mouse RAW 264.7 macrophages (Hadkar and Skidgel, 2001).

We have identified 4 more endogenous HEK293 proteins that are down-regulated in the presence of overexpressed WT MARCH1. CD112 and NKCC1 are expressed in dendritic cells, suggesting that MARCH1 may regulate cell shape during maturation and migration (Barbaro et al., 2017; Demian et al., 2019; Haghayegh Jahromi et al., 2024; Pende et al., 2006). Finally, two members of the solute carrier family 38 proteins, SNAT1 and SNAT2, were identified. The latter was further characterized since it showed marked cell surface downregulation in the presence of MARCH1 (Fig. 5B). SNAT2 is a neutral amino acid transporter acting at the plasma membrane. Homeostasis of the intracellular amino acid pool is highly regulated. Cellular metabolic pathways feeding into the tricyclic acid cycle can vary between cells, favoring either an increased glucose or glutamine metabolism as a source of carbon. In rapidly proliferating cells, such as T lymphocytes, increased glutamine transport is required to sustain metabolic needs (Carr et al., 2010). In cancer cells, glutamine is used as both a source of carbon and nitrogen (Yang et al., 2017). Following activation, rapid adaptation of amino acids transport in immune cells is key to the mounting of an effective response. Indeed, thrilling new results published recently demonstrated that in the tumor environment, dendritic cells depend on SNAT2 to acquire glutamine and initiate anti-cancer immunity (Guo et al., 2023). Thus, we can hypothesize that the drop in MARCH1 expression, which is associated with the maturation of DCs, is a prerequisite for the upregulation of SNAT2 and the acquisition of glutamine. However, many other players are likely involved in the control of amino acids and ions transport in various physiological conditions. It was of the utmost interest that the group of Rotin recently described that a bait, consisting of the SLC7A5 subunit of the LAT1 leucine transporter coupled to BioID, fished out SNAT2 and NKCC1 as proximity partners (Demian et al., 2019). As LAT1 is formed by the association between SLC7A5 and CD98, our results point to a role of MARCH1 in regulating the whole complex at different levels, and possibly in controlling the downstream activation of mTORC1 (Demian et al., 2019).

Altogether, our results point to a role of MARCH1 in cellular metabolism, which is subjected to a tight control in immune cells. Inflammation, antigen presentation and clonal expansion all command rapid changes in metabolism. Glutamine is a highly craved amino acid in activated dendritic cells, macrophages, B cells and T lymphocytes due to its role in protein synthesis and oxidative phosphorylation. To better comprehend the full spectrum of action of MARCH1 on the cell biology, we will need a better knowledge of its local interactome. This is a challenging task because of its low expression and the transient nature of the interactions with its targets. As part of our analysis, we have applied some arbitrary thresholds and filters that could be revisited when mining our data for new proteins that encounter MARCH1. For example, we have excluded proteins that were not found in MARCH1-transfected cells, but some were nevertheless relatively abundant with MARCH1W104A. SLC29A1 was one of them. This integral membrane protein is an important “equilibrative nucleoside transporter” that can also mediate uptake of anti-cancer nucleoside analogues, such as cytarabine (Megias-Vericat et al., 2022). In the future, the use of alternative, complementary BioID tools and more cell types will help broaden the MARCH proteins interaction networks.

### Author contributions

M.R., S.P., J.F.G. and J.T. designed the BioID constructs. R.B., W.M., D.G.R. and J.T. designed the experiments. R.B., W.M., M.A.M., L.C.Z.P, A.B.V. performed the experiments. D.F. performed the mass spectrometry analysis and wrote the associated protocol. R.B., W.M. and J.T. interpreted the data and wrote the manuscript. All authors revised and approved the final version of the paper.

## Acknowledgements

We thank Dr. Armelle Le Campion for helping with flow cytometry and Dr. Etienne Coyaud for useful discussions. J.T. holds the Saputo Research Chair. R.B. received scholarships from Diabète Québec and from the Faculty of Medicine of the University of Montreal. W.M received scholarships from the Natural Sciences and Engineering Research Council of Canada (NSERC).

---

**Table 1.**
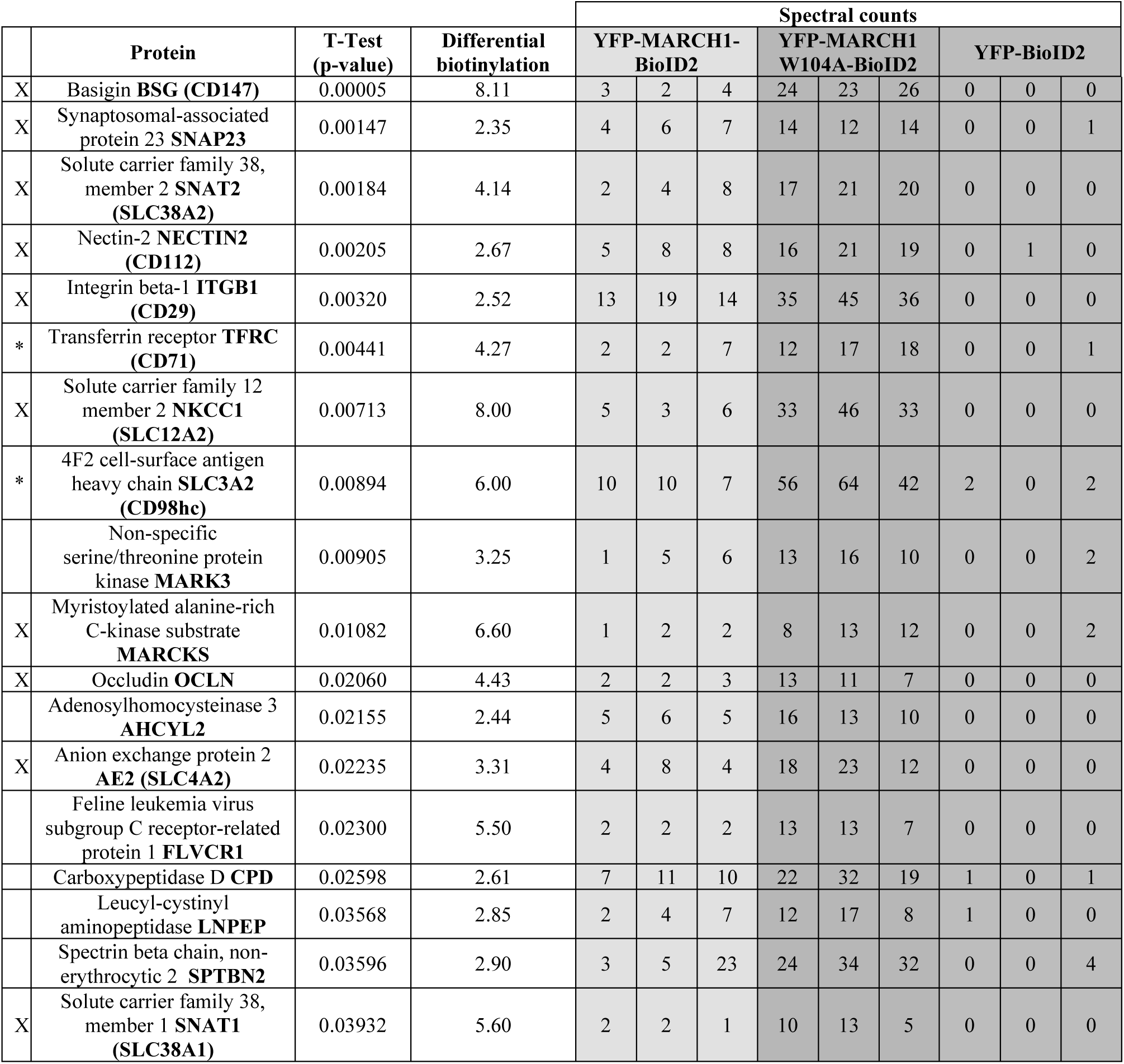
Potential MARCH1 targets. The proteins identified by mass spectrometry were filtered as described in the text and ranked in order of p-value. Differential biotinylation was calculated based on the ratio of spectral counts for YFP-MARCH1W104A-BioID2 over YFP-MARCH1-BioID2 and only the targets with a fold change > 2 were kept. Known targets of MARCH1 are identified by an asterisk in the left column (*), while an “X” indicates the proteins tested by flow cytometry (see figure 3) for their susceptibility to MARCH1-mediated down-regulation.

