## Supplemental figures for "A PROXIMITY LIGATION SCREEN IDENTIFIES SNAT2 AS A NOVEL TARGET OF THE MARCH1 E3 UBIQUITIN LIGASE"

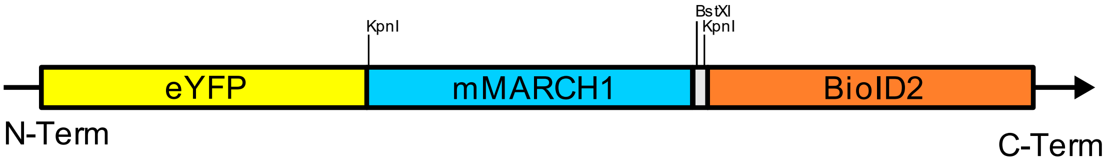


GCGGCCGCGCCACCATGGTTTCTAAAGGCGAGGAACTGTTCACCGGCGTGGTGCCTATTCTG
GTGGAACTGGATGGCGACGTGAACGGCCACAAGTTCTCCGTTTCTGGCGAAGGCGAAGGGG
ACGCCACATACGGAAAGCTGACCCTGAAGTTCATCTGCACCACCGGCAAGCTGCCTGTGCCT
TGGCCTACACTGGTCACCACATTTGGCTACGGCCTGCAGTGCTTCGCTAGATACCCCGACCAT
ATGAAGCTGCACGACTTCTTCAAGAGCGCCATGCCTGAGGGCTACGTGCAAGAGAGAACCAT
CTTCTTTAAGGACGACGGCAACTACAAGACCAGGGCCGAAGTGAAGTTCGAGGGCGACACC
CTGGTCAACAGAATCGAGCTGAAGGGCATCGACTTCAAAGAGGACGGCAACATCCTGGGCC
ACAAGCTCGAGTACAACTACAACAGCCACAACGTGTACATCATGGCCGACAAGCAGAAAAAC
GGCATCAAAGTGAACTTCAAGATCCGGCACAACATCGAGGACGGCTCTGTGCAGCTGGCCGA
TCACTACCAGCAGAACACACCTATCGGCGACGGACCTGTGCTGCTGCCTGATAACCACTACCT
GAGCTACCAGAGCGCCCTGAGCAAGGACCCTAACGAGAAGAGGGACCACATGGTGCTGCTG
GAATTCGTGACAGCCGCTGGCATCACACTCGGCATGGACGAGCTGTACAAGGGTACCAACCT
GACCATGAGCAACATGACCAGCAGCCACATCTGCTGCAACTTTCTGAACATGTGGAAGAAGT
CCAAGATCTCCACCATGTACTACCTGAACCAGGACGCCAAGCTGAGCAACCTGTTCCTGCAA
GCCAGCTCTCCCACAACCGGCACAGCTCCTAGAAGCCAGAGCAGACTGAGCGTGTGCCCTAG
CACACAGGACATCTGCAGAATCTGCCACTGTGAAGGCGACGAGGAAAGCCCTCTGATCACCC
CTTGCAGATGCACCGGCACACTGAGATTCGTGCACCAGAGCTGTCTGCACCAGTGGATCAAG
AGCAGCGACACCAGATGCTGCGAGCTGTGCAAATACGACTTCATCATGGAAACGAAGCTGAA
GCCCCTGAGGAAGTGGGAGAAGCTGCAGATGACCACCAGCGAGAGAAGAAAGATCTTCTGC
AGCGTGACCTTCCACGTGATCGCCGTGACCTGTGTCGTCTGGTCCCTGTACGTGCTGATCGAC
AGAACCGCCGAAGAGATCAAGCAGGGCGTGCTGGAATGGCCCTTCTGGACAAAACTGGTGG
TGGTGGCCATCGGCTTCACAGGCGGACTGGTGTTTATGTACGTGCAGTGCAAGGTGTACGTCC
AGCTGTGGCGGAGACTGAAGGCCTACAACAGAGTGATCTTCGTGCAGAACTGCCCCGACACC
GCTAACAAGCTGGAAAAGAACTTCCCCTGCAACGTGAACACCGAGATCAAGGACGCTGTGG
TGGTGCCAGTGCCTCAGACCGGATCTAACACACTGCCTACAGCTGAGGGCGCTCCTCCTGAA
GTGATCCCTGTTCCATGTACCTGGGGCAGCGCTAGCTTCAAGAACCTGATCTGGCTGAAAGAG
GTGGACAGCACCCAAGAGCGACTGAAAGAATGGAACGTGTCCTACGGCACAGCCCTGGTGG
CCGACAGACAAACAAAAGGCAGAGGCGGCCTGGGCAGAAAGTGGCTGTCTCAAGAAGGCG
GGCTGTACTTCTCCTTCCTGCTGAACCCCAAAGAGTTCGAGAACCTGCTGCAGCTGCCTCTGG
TTCTGGGCCTGTCTGTGTCTGAAGCCCTGGAAGAGATTACAGAGATCCCCTTCAGCCTGAAGT

GGCCCAACGACGTGTACTTCCAAGAGAAAAAGGTGTCCGGCGTGCTGTGCGAACTGTCCAA
GGACAAGCTGATCGTCGGCATCGGCATTAACGTGAACCAGAGAGAGATCCCAGAGGAAATCA
AGGACAGAGCCACCACACTGTACGAGATCACAGGCAAGGACTGGGACAGAAAAGAAGTGCT
GCTGAAGGTCCTGAAGCGGATCAGCGAGAATCTGAAGAAGTTCAAAGAGAAGTCCTTTAAA
GAGTTCAAGGGCAAGATCGAGAGCAAGATGCTGTACCTCGGCGAGGAAGTGAAGCTGCTCG
GCGAAGGCAAGATCACCGGAAAGCTCGTGGGCCTGAGCGAAAAAGGCGGCGCTCTGATCCT
GACCGAGGAAGGGATCAAAGAGATCCTGAGCGGCGAGTTCTCCCTGAGAAGAAGCTG*

**Figure S1: Map and nuclotide sequence of eYFP-MARCH1-BioID** YFP, MARCH1 and BioID domains of the fusion protein aswell as important restriction sites are mapped. Map colors correspond to highlight color in the nucleotide sequence.

atgaagaaggccgaaatgggacgattcagtatttccccggatgaagacagcagcagctacagttccaacagcgacttcaactactcctaccccaccaag
caagctgctctgaaaagccattatgcagatgtagatcctgaaaaccagaactttttacttgaatcgaatttggggaagaagaagtatgaaacagaatttcatc
caggtactacttcctttggaatgtcagtatttaatctgagcaatgcgattgtgggcagtggaatccttgggctttcttatgccatggctaatactggaattgctct
ttttataattctcttgacatttgtgtcaatattttccctgtattctgttcatctccttttgaagactgccaatgaaggagggtctttattatatgaacaattgggatataa
ggcatttggattagttggaaagcttgcagcatctggatcaattacaatgcagaacattggagctatgtcaagctacctcttcatagtgaaatatgagttgccttt
ggtgatccaggcattaacgaacattgaagataaaactggattgtggtatctgaacgggaactatttggttctgttggtgtcattggtggtcattcttcctttgtcg
ctgtttagaaatttaggatatttgggatataccagtggcctttccttgttgtgtatggtgttctttctgattgtggtcatttgcaagaaatttcaggttccgtgtcctgt
ggaagctgctttgataattaacgaaacaataaacaccaccttaacacagccaacagctcttgtacctgctttgtcacataacgtgactgaaaatgactcttgc
agacctcactattttattttcaactcacagactgtctatgctgtgccaattctgatcttttcatttgtctgtcatcctgctgttcttcccatctatgaagaactgaaag
accgcagccgtagaagaatgatgaatgtgtccaagatttcatttttgctatgtttctcatgtatctgcttgccgccctctttggatacctaacattttacgaacat
gttgagtcagaattgcttcatacctactcttctatcttgggaactgatattcttcttctcattgtccgtctggctgtgttaatggctgtgaccctgacagtaccagta
gttattttcccaatccggagttctgtaactcacttgttgtgtgcatcaaaagatttcagttggtggcgtcatagtctcattacagtgtctatcttggcatttaccaat
ttacttgtcatctttgtcccaactattagggatatctttggttttattggtgcatctgcagcttctatgttgatttttattcttccttctgccttctatatcaagttggtgaa
gaaagaacctatgaaatctgtacaaaagattggggctttgttcttcctgttaagtggtgtactggtgatgaccggaagcatggccttgattgttttggattgggt
acacaatgcacctggaggtggccatGACTACAAGGACGACGACGACAAG

**Figure S2: Nucleotide sequence of SNAT2-FLAG.** Lowercase letters constitute the sequence of SNAT2 while uppercase letters represent the sequence of the FLAG-TAG.

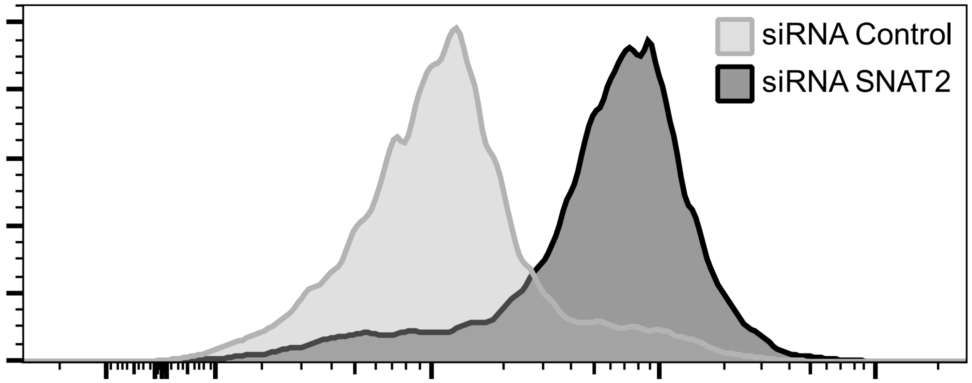


**Figure S3: siRNA confirmation of the specificity of the SNAT2 antibody.** HEK293 cells were transfected with SNAT2 specific and control siRNAs as described above. Anti SNAT2 antibody was used to detect intracellular SNAT2 in both siRNA condition by flow cytometry.

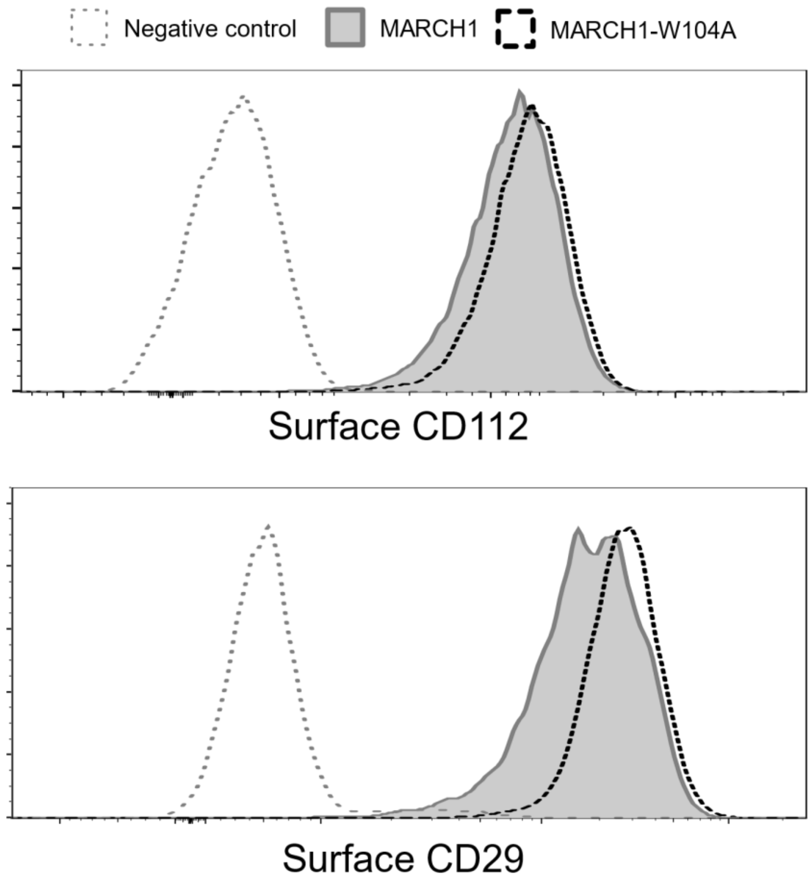


**Figure S4: The presence of MARCH1 but not MARCH1W104A has an effect on the surface expression of CD29 and CD112.** HEK293 cells were transfected with either MARCH1 or MARCH1W104A. Levels of surface endogenous CD29 and CD112 were measured by flow-cytometry after 48 hours. Negative controls consist of a secondary antibody staining for CD112 and an ia

**Table S1: Conformation and known membrane association of proteins identified in MARCH1 BioID experiments.**

| Protein | Trans membrane | Plasma membrane association | Cytoplasmic lysine residue | References | Notes |
|---|---|---|---|---|---|
| Basigin **BSG (CD147)** | Yes | Yes | Yes | (Kanekura, Chen and Kanzaki, 2002) | |
| Synaptosomal-associated protein 23 **SNAP23** | No | Yes | Yes | (Hatsuzawa and Sakurai, 2020; Naskar and Puri, 2017) | |
| Solute carrier family 38, member 2 **SNAT2 (SLC38A2)** | Yes | Yes | Yes | (Zhang, Zander and Grewer, 2011) | |
| Nectin-2 **NECTIN2 (CD112)** | Yes | Yes | Yes | (Samanta et al., 2012) | |
| Integrin beta-1 **ITGB1 (CD29)** | Yes | Yes | Yes | (Zuidema et al., 2022) | |
| Transferrin receptor **TFRC (CD71)** | Yes | Yes | Yes | (Jabara et al., 2016) | |
| Solute carrier family 12 member 2 **NKCC1 (SLC12A2)** | Yes | Yes | Yes | (Payne et al., 1995; Yang, Wang and Cao, 2020) | |
| 4F2 cell-surface antigen heavy chain **SLC3A2 (CD98hc)** | Yes | Yes | Yes | (Quackenbush et al., 1987) | |
| Non-specific serine/threonine protein kinase **MARK3** | No | Yes | Yes | (Moravcevic et al., 2010) | Associated to phospholipids |
| Myristoylated alanine-rich C-kinase substrate **MARCKS** | No | Yes | Yes | (Carballo et al., 1999; Issara-Amphorn et al., 2023; Thelen et al., 1991) | Associated to PKC |
| Occludin **OCLN** | Yes | Yes | Yes | (Elias et al., 2009) | |
| Adenosyl homocysteinase 3 **AHCYL2 (Long-IRBIT)** | No | Possible | Yes | (Ando, Mizutani and Mikoshiba, 2009) | (Uniprot) Note: Associates with membranes when phosphorylated, probably through interaction with ITPR1. |
| Anion exchange protein 2 **AE2 (SLC4A2)** | Yes | Yes | Yes | (Aranda et al., 2004; Itoh et al., 2021) | Associates with long-IRBIT for lysosomal degradation. |
| Feline leukemia virus subgroup C receptor-related protein 1 **FLVCR1** | Yes | Yes | Yes | (Fiorito and Tolosano, 2024; Quigley et al., 2004; Ri et al., 2024) | Associated to heme and choline transport. |
| Carboxypeptidase D **CPD** | Yes | Yes | Yes | (Kuroki et al., 1995) | |
| Leucyl-cystinyl aminopeptidase **LNPEP** | Yes | Yes | Yes | (Rogi et al., 1996) | (Uniprot) In brain only the membrane-bound form is found. The protein resides in intracellular vesicles |

| | | | | | |
|---|---|---|---|---|---|
| | | | | | together with GLUT4 and can then translocate to the cell surface in response to insulin and/or oxytocin. |
| Spectrin beta chain, non-erythrocytic 2 **SPTBN2** | No | Possible | Yes | (Deng et al., 2024) | SPTBN2 suppresses ferroptosis in NSCLC cells by facilitating SLC7A11 membrane trafficking and localization |
| Solute carrier family 38, member 1 **SNAT1 (SLC38A1)** | Yes | Yes | Yes | (Melone et al., 2004) | |

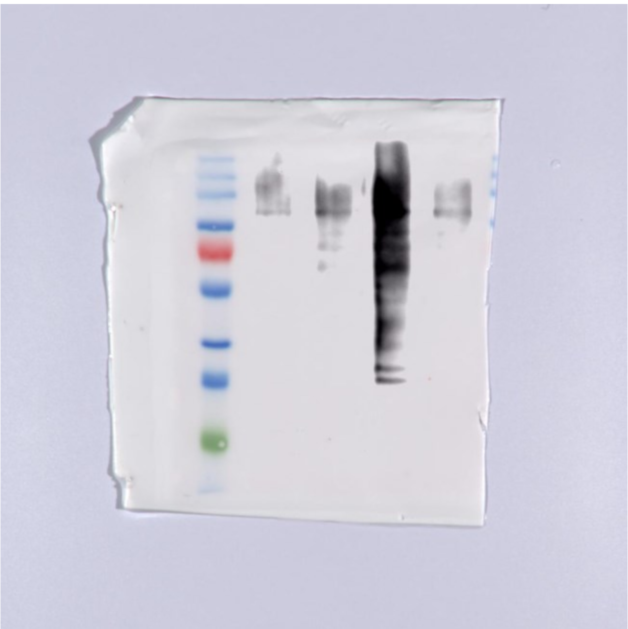

**Figure S5: Blot transparency related to figure 5C**

**Refererences:**


**Ando, H., Mizutani, A. and Mikoshiba, K.** (2009). An IRBIT homologue lacks binding activity to inositol 1,4,5-trisphosphate receptor due to the unique N-terminal appendage. *J Neurochem* **109**, 539-50.

**Aranda, V., Martinez, I., Melero, S., Lecanda, J., Banales, J. M., Prieto, J. and Medina, J. F.** (2004). Shared apical sorting of anion exchanger isoforms AE2a, AE2b1, and AE2b2 in primary hepatocytes. *Biochem Biophys Res Commun* **319**, 1040-6.

**Carballo, E., Pitterle, D. M., Stumpo, D. J., Sperling, R. T. and Blackshear, P. J.** (1999). Phagocytic and macropinocytic activity in MARCKS-deficient macrophages and fibroblasts. *Am J Physiol* **277**, C163-73.

**Deng, J., Lin, X., Qin, J., Li, Q., Zhang, Y., Zhang, Q., Ji, C., Shen, S., Li, Y., Zhang, B. and Lin, N.** (2024). SPTBN2 suppresses ferroptosis in NSCLC cells by facilitating SLC7A11 membrane trafficking and localization. *Redox Biol* **70**, 103039.

**Elias, B. C., Suzuki, T., Seth, A., Giorgianni, F., Kale, G., Shen, L., Turner, J. R., Naren, A., Desiderio, D. M. and Rao, R.** (2009). Phosphorylation of Tyr-398 and Tyr-402 in occludin prevents its interaction with ZO-1 and destabilizes its assembly at the tight junctions. *J Biol Chem* **284**, 1559-69.

**Fiorito, V. and Tolosano, E.** (2024). Unearthing FLVCR1a: tracing the path to a vital cellular transporter. *Cell Mol Life Sci* **81**, 166.

**Hatsuzawa, K. and Sakurai, C.** (2020). Regulatory Mechanism of SNAP23 in Phagosome Formation and Maturation. *Yonago Acta Medica* **63**, 135-145.

**Issara-Amphorn, J., Sjoelund, V. H., Smelkinson, M., Montalvo, S., Yoon, S. H., Manes, N. P. and Nita-Lazar, A.** (2023). Myristoylated, alanine-rich C-kinase substrate (MARCKS) regulates toll-like receptor 4 signaling in macrophages. *Scientific Reports* **13**, 19562.

**Itoh, R., Hatano, N., Murakami, M., Mitsumori, K., Kawasaki, S., Wakagi, T., Kanzaki, Y., Kojima, H., Kawaai, K., Mikoshiba, K. et al.** (2021). Both IRBIT and long-IRBIT bind to and coordinately regulate Cl(-)/HCO(3)(-) exchanger AE2 activity through modulating the lysosomal degradation of AE2. *Scientific Reports* **11**, 5990.

**Jabara, H. H., Boyden, S. E., Chou, J., Ramesh, N., Massaad, M. J., Benson, H., Bainter, W., Fraulino, D., Rahimov, F., Sieff, C. et al.** (2016). A missense mutation in TFRC, encoding transferrin receptor 1, causes combined immunodeficiency. *Nat Genet* **48**, 74-8.

**Kanekura, T., Chen, X. and Kanzaki, T.** (2002). Basigin (CD147) is expressed on melanoma cells and induces tumor cell invasion by stimulating production of matrix metalloproteinases by fibroblasts. *Int J Cancer* **99**, 520-8.

**Kuroki, K., Eng, F., Ishikawa, T., Turck, C., Harada, F. and Ganem, D.** (1995). gp180, a host cell glycoprotein that binds duck hepatitis B virus particles, is encoded by a member of the carboxypeptidase gene family. *J Biol Chem* **270**, 15022-8.

**Melone, M., Quagliano, F., Barbaresi, P., Varoqui, H., Erickson, J. D. and Conti, F.** (2004). Localization of the glutamine transporter SNAT1 in rat cerebral cortex and neighboring structures, with a note on its localization in human cortex. *Cereb Cortex* **14**, 562-74.

**Moravcevic, K., Mendrola, J. M., Schmitz, K. R., Wang, Y. H., Slochower, D., Janmey, P. A. and Lemmon, M. A.** (2010). Kinase associated-1 domains drive MARK/PAR1 kinases to membrane targets by binding acidic phospholipids. *Cell* **143**, 966-77.

**Naskar, P. and Puri, N.** (2017). Phosphorylation of SNAP-23 regulates its dynamic membrane association during mast cell exocytosis. *Biol Open* **6**, 1257-1269.

**Payne, J. A., Xu, J. C., Haas, M., Lytle, C. Y., Ward, D. and Forbush, B., 3rd.** (1995). Primary structure, functional expression, and chromosomal localization of the bumetanide-sensitive Na-K-Cl cotransporter in human colon. *J Biol Chem* **270**, 17977-85.

**Quackenbush, E., Clabby, M., Gottesdiener, K. M., Barbosa, J., Jones, N. H., Strominger, J. L., Speck, S. and Leiden, J. M.** (1987). Molecular cloning of complementary DNAs encoding the heavy chain of the human 4F2 cell-surface antigen: a type II membrane glycoprotein involved in normal and neoplastic cell growth. *Proceedings of the National Academy of Sciences* **84**, 6526-6530.

**Quigley, J. G., Yang, Z., Worthington, M. T., Phillips, J. D., Sabo, K. M., Sabath, D. E., Berg, C. L., Sassa, S., Wood, B. L. and Abkowitz, J. L.** (2004). Identification of a human heme exporter that is essential for erythropoiesis. *Cell* **118**, 757-66.

**Ri, K., Weng, T. H., Claveras Cabezudo, A., Josting, W., Zhang, Y., Bazzone, A., Leong, N. C. P., Welsch, S., Doty, R. T., Gursu, G. et al.** (2024). Molecular mechanism of choline and ethanolamine transport in humans. *Nature* **630**, 501-508.

**Rogi, T., Tsujimoto, M., Nakazato, H., Mizutani, S. and Tomoda, Y.** (1996). Human placental leucine aminopeptidase/oxytocinase. A new member of type II membrane-spanning zinc metallopeptidase family. *J Biol Chem* **271**, 56-61.

**Samanta, D., Ramagopal, U. A., Rubinstein, R., Vigdorovich, V., Nathenson, S. G. and Almo, S. C.** (2012). Structure of Nectin-2 reveals determinants of homophilic and heterophilic interactions that control cell-cell adhesion. *Proc Natl Acad Sci U S A* **109**, 14836-40.

**Thelen, M., Rosen, A., Nairn, A. C. and Aderem, A.** (1991). Regulation by phosphorylation of reversible association of a myristoylated protein kinase C substrate with the plasma membrane. *Nature* **351**, 320-322.

**Yang, X., Wang, Q. and Cao, E.** (2020). Structure of the human cation-chloride cotransporter NKCC1 determined by single-particle electron cryo-microscopy. *Nat Commun* **11**, 1016.

**Zhang, Z., Zander, C. B. and Grewer, C.** (2011). The C-terminal domain of the neutral amino acid transporter SNAT2 regulates transport activity through voltage-dependent processes. *Biochem J* **434**, 287-96.

**Zuidema, A., Atherton, P., Kreft, M., Hoekman, L., Bleijerveld, O. B., Nagaraj, N., Chen, N., Fassler, R. and Sonnenberg, A.** (2022). PEAK1 Y635 phosphorylation regulates cell migration through association with Tensin3 and integrins. *J Cell Biol* **221**.
